# Bacterial community patterns of the Antarctic stalked jellyfish *Haliclystus antarcticus* and antimicrobial activity from a gut-associated *Oceanisphaera ostreae* isolate

**DOI:** 10.64898/2026.11.20.768063

**Authors:** Federica Montesanto, Joseph B. Mangun, Grayce E. Dyer, Jaime R. Cabrera-Pardo, Sandra Loesgen

## Abstract

Host-associated microbiomes are increasingly recognized as important components of cnidarian biology, yet their diversity and organization remain poorly understood in non-coral lineages, particularly in polar environments. Staurozoans, or stalked jellyfishes, are a small and understudied group of benthic medusozoans whose associated bacterial communities remain largely unexplored. Here, we used 16S rRNA gene amplicon sequencing to characterize bacterial communities associated with the Antarctic stalked jellyfish *Haliclystus antarcticus* collected from two coastal sites in the South Shetland Islands, Byers Bay and Risopatron. We examined three anatomical compartments (gut, peduncle, and tentacles) to investigate patterns of bacterial community diversity and composition. Gut samples exhibited the greatest variability in richness-based alpha diversity, whereas peduncle and tentacle samples displayed more constrained diversity profiles. Across compartments, 47 amplicon sequence variants (ASVs) were shared, indicating a recurrent bacterial component within the host-associated communities. Gut samples exhibited the strongest differential-abundance signal, while genera such as *Polaribacter* and *Planktotalea* were identified as indicators of tentacle-associated communities. In parallel, we isolated and characterized a gut-associated bacterium identified as *Oceanisphaera ostreae*. Genomic and metabolomic analyses revealed diverse biosynthetic potential and multiple metabolite-related features. Preliminary bioactivity screening indicated antibacterial activity against *Staphylococcus aureus*, with no cytotoxicity detected against HCT-116 cells under the tested conditions. Together, these results provide the first characterization of the *H. antarcticus* microbiome, demonstrating anatomical and geographic variation in bacterial communities and identifying a cultivable bacterial associate with potentially relevant metabolic capabilities.

**IMPORTANCE:** Understanding how marine invertebrates associate with microorganisms is essential for revealing the complexity of host–microbe relationships, particularly in extreme and poorly explored environments such as Antarctica. By investigating the Antarctic stalked jellyfish *Haliclystus antarcticus*, this study extends microbiome research to a largely overlooked cnidarian lineage and provides evidence that bacterial associations are not uniform across the host body or geographic locations. Beyond describing community composition, the characterization of a cultivable gut-associated *Oceanisphaera ostreae* strain highlights the potential of understudied host-associated bacteria as sources of metabolic diversity and bioactive compounds. By combining community profiling with genomic, metabolomic, and preliminary bioactivity analyses, this work establishes a foundation for investigating the ecological significance and functional potential of bacterial associations in polar cnidarians.

## INTRODUCTION

In cnidarians, bacterial communities have been studied most extensively in anthozoans, particularly reef-building corals, where host-associated microorganisms contribute to nutrient cycling, carbon and nitrogen metabolism, and host health (1–3). However, several cnidarian lineages remain microbiologically underexplored, limiting our understanding of how host identity, anatomy, ecology, and environmental conditions influence bacterial community structure across Cnidaria (4–10). Staurozoa, or stalked jellyfishes, remain microbiologically understudied, with the first bacterial microbiome characterization reported only recently for *Haliclystus auricula* James-Clark, 1863 (7).

Unlike most medusozoans, staurozoans have a benthic adult stage that remains attached to the substrate rather than swimming freely. Their body plan combines polyp-like and medusa-like characteristics, with an aboral peduncle bearing an adhesive basal disc and an oral region comprising the calyx, manubrium, arms, and secondary tentacles (11, 12). These anatomical regions perform different functions and are exposed to distinct environmental and physiological conditions. The peduncle provides attachment to the substrate, the tentacles mediate prey capture and contact with the surrounding environment, and the gastrovascular cavity is involved in digestion. Studies on free-swimming jellyfish have demonstrated that bacterial community composition can vary among anatomical regions, including the exumbrella, oral arms, and gastric cavity of *Aurelia solida* Browne, 1905 (13), as well as different tissues of *Rhopilema nomadica* Galil, 1990 (14). These findings raise the possibility that anatomical compartmentalization also contributes to bacterial community structure in staurozoans, although this remains largely unexplored.

The Antarctic stalked jellyfish *Haliclystus antarcticus* Pfeffer, 1889 provides an opportunity to investigate bacterial associations in a cnidarian adapted to cold-water benthic environments. Previous studies have primarily addressed its taxonomy, morphology, internal anatomy, life cycle, and trophic ecology. Taxonomic and anatomical investigations have revealed considerable morphological complexity, including specialized structures associated with attachment, feeding, and reproduction (12, 15). In staurozoans, the peduncle is the structure directly involved in attachment to the substrate, the tentacular region is involved in prey capture and environmental contact, and the gastrovascular/gut region is linked to digestion and internal processing of food-associated material (11, 12).

Ecologically, stalked jellyfishes are cryptic benthic predators that often occur attached to algae, rocks, or other hard substrates in intertidal and shallow subtidal habitats. Available observations indicate that stauromedusae feed on both planktonic and benthic prey, including copepods, amphipods, and other small invertebrates, and may themselves be consumed by higher trophic levels. Such trophic interactions are relevant for microbiome assembly because the gut and feeding structures may receive bacteria from prey, suspended particles, surrounding seawater, and the benthic microhabitat. In *H. antarcticus* from South Georgia, Davenport reported both planktonic and benthic prey in gut contents, while diet studies in related *Haliclystus* species also indicate size- and prey-dependent feeding patterns (16, 17).

These feeding habits may expose the gastrovascular cavity to bacteria originating from prey and ingested environmental material, whereas external structures remain in direct contact with surrounding seawater and benthic substrates. Such differences in exposure provide a biological rationale for investigating whether bacterial communities vary among anatomical compartments.

Despite increasing interest in cnidarian-associated microbiomes, bacterial communities associated with Staurozoa remain poorly characterized. The first microbiome study of a stalked jellyfish, focused on *Haliclystus auricula* James-Clark, 1863 from the Barents Sea, showed that staurozoans can host diverse bacterial assemblages dominated by cold-adapted taxa, including genera such as *Colwellia* and *Marinomonas* (7). The microbiome of the Antarctic species *H. antarcticus* remains uncharacterized, thus investigating this species provides an opportunity to extend current knowledge of bacterial associations in polar cnidarians and assess the relative contributions of anatomical compartment and sampling location to microbiome variation. Cultivation-based approaches can further complement community profiling by enabling the genomic and chemical characterization of individual host-associated bacterial isolates.

Here, we characterize the bacterial microbiome of *H. antarcticus* collected from two sites in the South Shetland Islands, Byers Bay and Risopatron, using 16S rRNA gene amplicon sequencing. We investigated three anatomical compartments (gut, peduncle, and tentacles) to determine whether bacterial community diversity and composition differ across the host body and between collection sites. Specifically, we aimed to: (i) characterize the diversity and taxonomic composition of host-associated bacterial communities; (ii) assess variation in bacterial assemblages among anatomical compartment and sampling site; (iii) identify shared and compartment-associated bacterial taxa; and (iv) complement community profiling with cultivation-based characterization of a gut-derived bacterial isolate through whole-genome analysis, genome mining, metabolite profiling, and preliminary bioactivity screening.

## RESULTS

### Host-associated bacterial diversity

We characterized bacterial communities associated with three anatomical compartments of *H. antarcticus* (gut, peduncle and tentacles) from individuals collected at Byers Bay and Risopatron in the South Shetland Islands (Fig. 1). After quality filtering, denoising, chimera removal, removal of non-bacterial, mitochondrial and chloroplast sequences, and exclusion of low-quality samples, the final dataset contained 565 bacterial ASVs across host-associated compartment samples.

**FIG 1.**
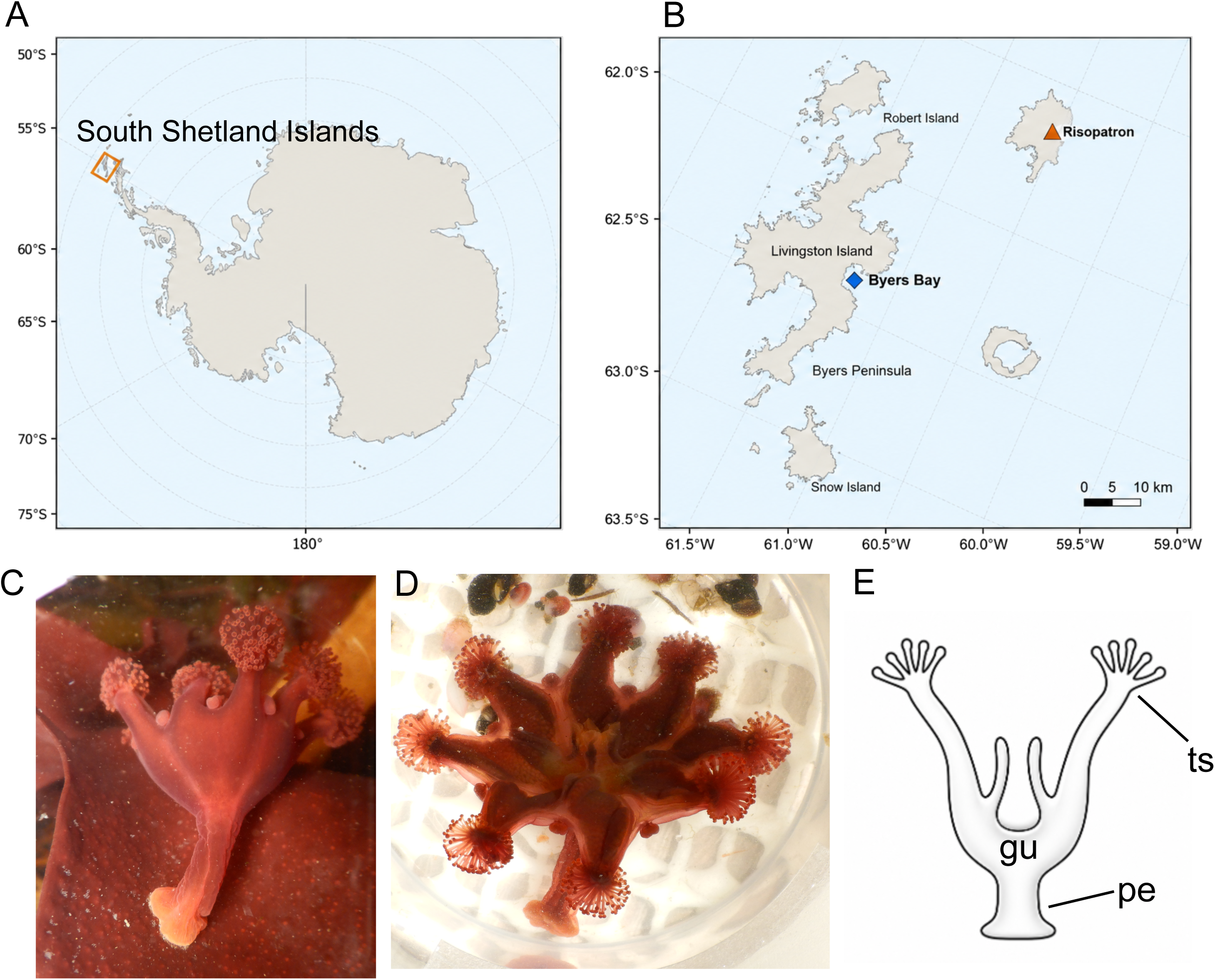
Sampling locations and anatomical overview of *Haliclystus antarcticus*. (A) Antarctic map showing the position of the South Shetland Islands. The orange box indicates the sampling region. (B) Detailed map of Livingston Island and surrounding islands showing the two sampling sites: Byers Bay and Risopatron. (C,D) Representative images of *H. antarcticus* individuals collected from the South Shetland Islands. (E) Schematic representation of the dissected host compartments used for 16S rRNA gene microbiome profiling: gut/gastrovascular cavity (*gu*), peduncle (*pe*), and tentacles/tentacular structures (*ts*).

Alpha diversity differed among host compartments, although variation within compartments was substantial (Fig. 2A). Gut samples showed the broadest spread in observed ASV richness and Fisher’s alpha, including several high-richness samples. Peduncle samples generally showed lower richness and lower dispersion, whereas tentacle samples had intermediate to high diversity depending on the metric. Shannon and inverse Simpson diversity showed less extreme separation among compartments than richness-based metrics, indicating that compartment differences were driven partly by differences in rare or low-abundance ASVs rather than only by dominant taxa.

**FIG 2.**
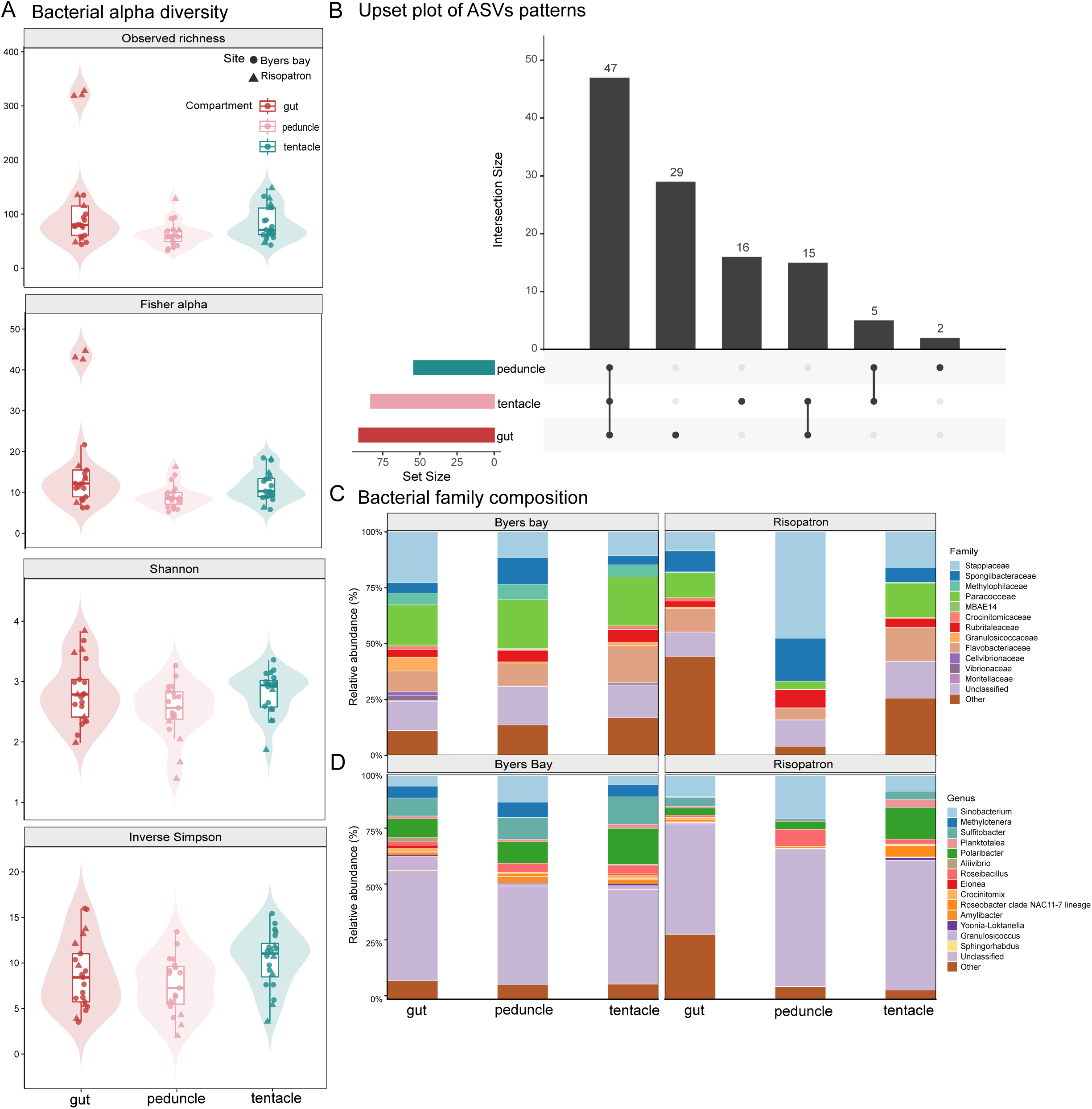
Diversity and taxonomic composition of bacterial communities associated with *H. antarcticus* compartments. (A) Alpha diversity of bacterial communities across gut, peduncle and tentacle samples, shown using observed ASV richness, Fisher’s alpha, Shannon diversity and inverse Simpson diversity. Violin plots show the distribution of values, boxplots show median and interquartile range, and points represent individual samples. (B) UpSet plot showing ASV sharing among gut, peduncle and tentacle samples. Bars indicate intersection size, and connected points indicate the compartment combinations represented by each intersection. (C,D) Relative abundance of dominant bacterial families (C) and genera (D) across host compartments and sampling sites. Lower-abundance taxa were grouped as “Other”.

Overall, these patterns suggest that bacterial diversity in *H. antarcticus* is not uniformly distributed across the body. The gut compartment was particularly variable, consistent with greater exposure to prey-associated, ingested, or transient bacterial taxa, whereas peduncle and tentacle samples showed more constrained diversity profiles.

Despite compartment-level variation in diversity, a substantial fraction of ASVs was shared among host compartments (Fig. 2B). The largest intersection consisted of 47 ASVs shared across gut, peduncle and tentacle samples, indicating the presence of a common host-associated bacterial pool. The other displayed disjoint intersections comprised 29 ASVs unique to the gut set, 16 unique to tentacles, 2 unique to peduncle, 15 shared by gut and tentacles only, and 5 shared by peduncle and tentacles only.

### Taxonomic composition and bacterial community structure

Family- and genus-level composition revealed both shared and compartment-specific bacterial patterns (Fig. 2C,D). At the family level, host-associated communities were composed primarily of bacterial groups including *Stappiaceae, Spongiibacteraceae, Methylophilaceae, Paracoccaceae, Flavobacteriaceae, Moritellaceae* and several unclassified bacterial lineages. The relative contribution of these families varied among gut, peduncle and tentacle samples and also differed between Byers Bay and Risopatron.

At the genus level, several taxa contributed prominently to the host-associated microbiome, including *Sinobacterium*, *Methylotenera*, *Sulfitobacter*, *Planktotalea*, *Polaribacter*, *Roseibacillus*, *Eionea*, *Aliivibrio* and members of the *Roseobacter* clade NAC11-7 lineage. A large fraction of reads remained unclassified at genus level, particularly in some site-compartment combinations, indicating that part of the *H. antarcticus*-associated bacterial community is not well resolved by current reference databases.

The taxonomic profiles suggest that *H. antarcticus* hosts a mixed bacterial assemblage containing common marine bacterial groups, cold-water-associated taxa and poorly classified lineages. The relative abundance of dominant families and genera differed across anatomical compartments, supporting the idea that host anatomy contributes to microbiome structure.

Ordination analyses showed that host-associated bacterial communities were structured by both sampling site and anatomical compartment (Fig. 3A,B). Bray–Curtis PCoA explained 21.6% of the variation on the first axis and 12.6% on the second axis. Weighted UniFrac PCoA explained 37.4% of the variation on the first axis and 17.9% on the second axis. In both ordinations, samples showed partial separation by compartment, with tentacle samples tending to occupy a distinct region relative to gut and peduncle samples. Sampling site also contributed to the ordination structure, particularly along the first ordination axes.

**FIG 3.**
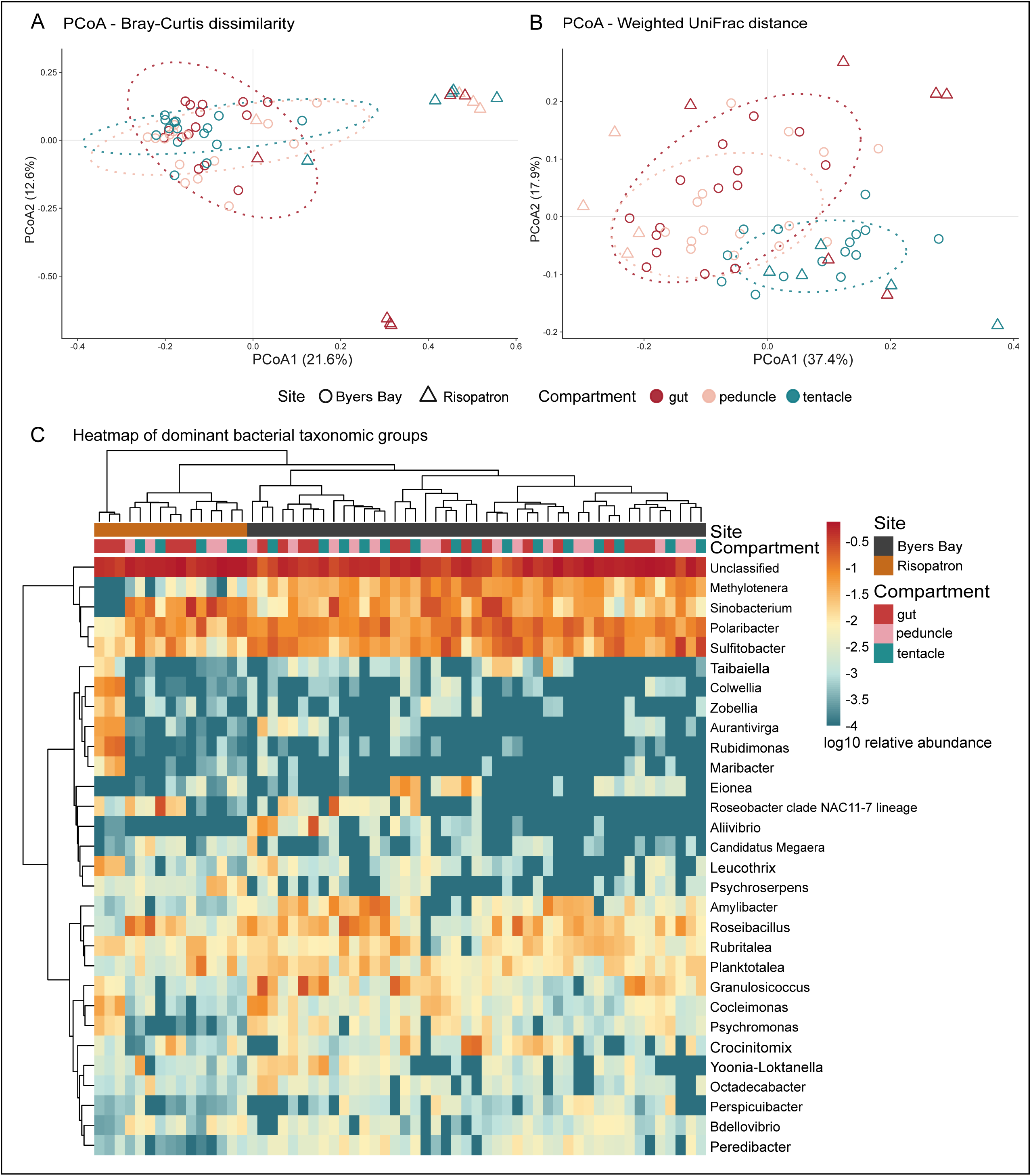
Beta diversity and dominant genus-level bacterial structure in *H. antarcticus*. (A) Principal coordinate analysis (PCoA) based on Bray–Curtis dissimilarity among host-associated bacterial communities. (B) PCoA based on weighted UniFrac distance. In both ordinations, points represent individual compartment samples, colours indicate host compartment, and shapes indicate sampling site. (C) Heatmap of dominant bacterial genera across host-associated samples. Values represent log10-transformed relative abundance. Rows and columns are hierarchically clustered, and sample annotations indicate sampling site and host compartment.

PERMANOVA confirmed significant effects of both site and compartment. For Bray–Curtis dissimilarity, site explained 16.9% of community variation (F = 12.65, *P* = 0.0001), whereas compartment explained 8.1% (F = 3.03, *P* = 0.0001). For weighted UniFrac distance, compartment explained a larger fraction of phylogenetic community variation (*R*² = 15.5%, F = 5.79, *P* = 0.0001), while site also remained significant (*R*² = 9.5%, F = 7.12, P = 0.0001). These results indicate that geographic context and host anatomical compartment both contribute to structuring the H. antarcticus microbiome.

A genus-level heatmap further supported these patterns (Fig. 3C). Samples clustered according to shifts in dominant bacterial genera, with several taxa showing compartment- or site-associated abundance patterns. Genera such as *Methylotenera*, *Sinobacterium*, *Polaribacter* and *Sulfitobacter* were among the more prominent taxa across samples, whereas taxa such as *Colwellia*, *Zobellia*, *Aurantivirga*, *Rubritalea*, *Maribacter*, *Aliivibrio* and others showed stronger enrichment in subsets of samples. This clustering suggests that community differentiation is driven not only by broad changes in diversity but also by consistent shifts in specific bacterial taxa.

### Compartment-associated bacterial taxa and co-abundance

Differential-abundance analysis using DESeq2 identified bacterial ASVs associated with anatomical compartments of *H. antarcticus* (Fig. 4A–C). In comparisons between gut and peduncle samples (Fig. 4a) and between gut and tentacle samples (Fig. 4B), the significant ASVs displayed were enriched in the gut. These included ASVs assigned to *Rubidimonas*, *Aurantivirga*, *Maribacter*, *Thiothrix*, *Zobellia*, *Rubritalea*, *Colwellia*, *Leucothrix*, *Tenacibaculum*, and *Taibaiella*, among others. In contrast, the comparison between peduncle and tentacle samples revealed differentially abundant ASVs enriched in both compartments, with several ASVs assigned to *Sulfitobacter*, *Eionea*, *Peredibacter*, *Lewinella*, *Rubidimonas*, *Colwellia*, and *Tenacibaculum* showing enrichment in tentacle samples (Fig. 4C).

**FIG 4.**
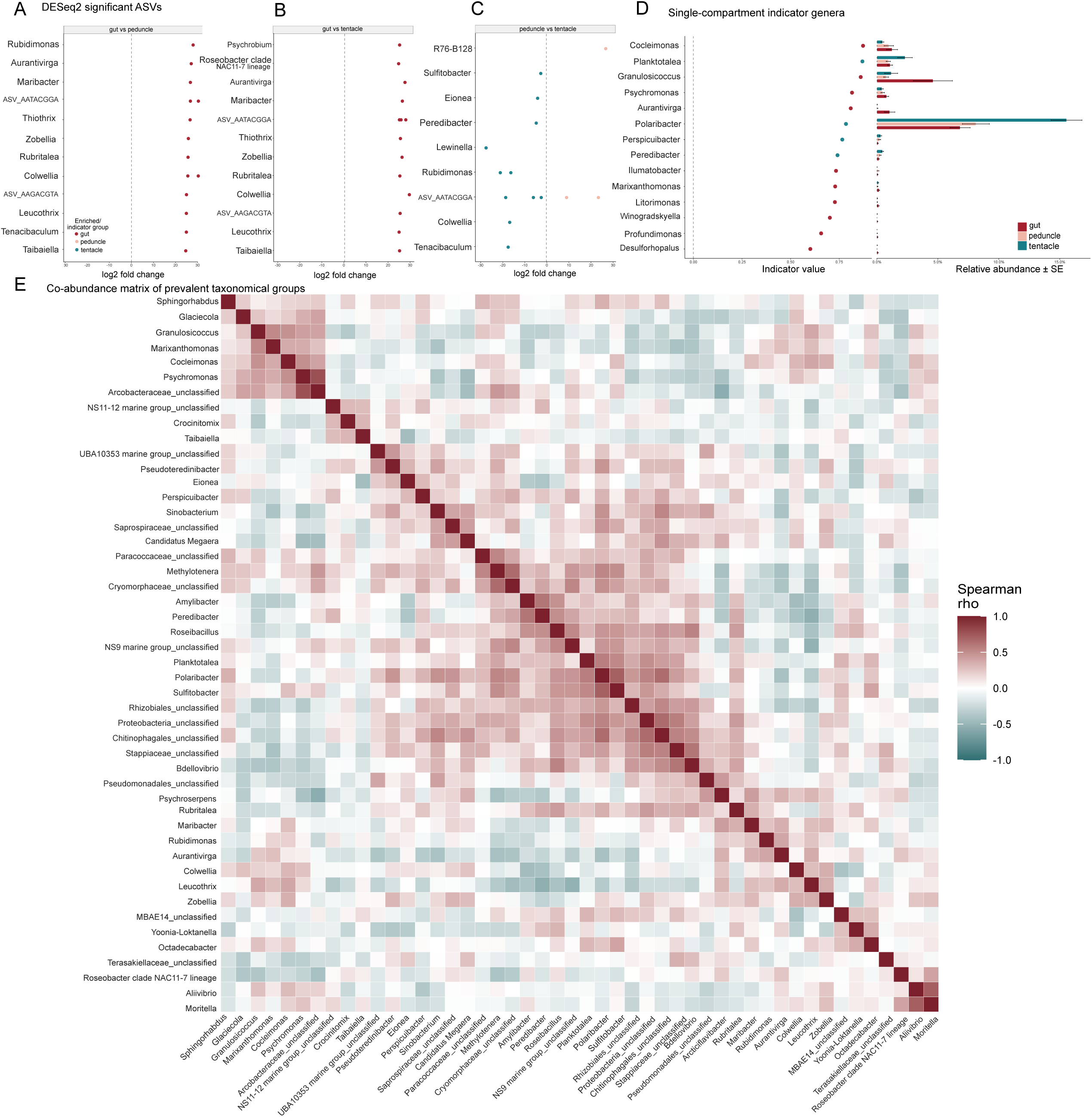
Differential abundance, indicator, and co-abundance patterns of bacterial taxa associated with anatomical compartments of *H. antarcticus*. (A–C) Pairwise differential-abundance analysis of bacterial amplicon sequence variants (ASVs) using DESeq2 for (A) gut versus peduncle, (B) gut versus tentacle, and (C) peduncle versus tentacle. Points represent significantly differentially abundant ASVs (Benjamini–Hochberg-adjusted *P* ≤ 0.05), labelled according to their lowest available taxonomic assignment. (D) Single-compartment indicator genera identified using indicator value analysis (IndVal). Points indicate indicator values for genera significantly associated with either gut or tentacle samples (*P* ≤ 0.05). Horizontal bars represent the mean relative abundance ± standard error of each genus across the three anatomical compartments. (E) Genus-level co-abundance matrix based on Spearman correlations calculated from centred log-ratio (CLR)-transformed relative abundance profiles of prevalent bacterial genera.

Complementary indicator value analysis (IndVal) identified significant single-compartment indicator genera associated with gut and tentacle samples, whereas no significant peduncle-specific indicators were detected under the applied criteria (Fig. 4D). Gut-associated indicators included *Cocleimonas*, *Granulosicoccus*, *Psychromonas*, *Aurantivirga*, *Ilumatobacter*, *Marixanthomonas*, *Litorimonas*, *Winogradskyella*, *Profundimonas*, and *Desulforhopalus*. Tentacle-associated indicator genera included *Planktotalea*, *Polaribacter*, *Perspicuibacter*, and *Peredibacter*. Among these, *Polaribacter* showed a relatively high abundance in tentacle samples, consistent with its identification as a tentacle-associated indicator.

Exploratory genus-level co-abundance analysis revealed structured positive and negative correlations among prevalent bacterial genera (Fig. 4E). The CLR–Spearman correlation matrix showed groups of positively correlated genera, including a cluster containing *Sphingorhabdus*, *Glaciecola*, *Granulosicoccus*, *Marixanthomonas*, *Cocleimonas*, and *Psychromonas*, and another containing genera such as *Planktotalea*, *Polaribacter*, *Sulfitobacter*, and *Peredibacter*. Negative correlations were also observed between several groups of genera, indicating contrasting abundance patterns across samples.

Together, these analyses revealed distinct compartment-associated taxonomic patterns, with gut and tentacle samples exhibiting different sets of enriched and indicator bacterial taxa.

### Cultivation and genome-based identification of *Oceanisphaera ostreae*

A bacterial strain (designated FM48a) was recovered from intestinal material of *H. antarcticus* specimen. The FM48a draft genome was 3,565,580 bp in seven contigs, with an N50 of 1,016,169 bp and GC content of 49.321%. The genome annotation reported 3,221 genes, and the sequencing summary reported 109x raw coverage and 95x coverage for the primary genome assembly. Reciprocal FastANI identified *O. ostreae* CCUG 60525, an assembly derived from type material, as the closest genome to FM48a. Mean reciprocal ANI was 98.5815%, with a minimum reciprocal alignment fraction of 0.8835, exceeding the species-level screening criteria used here. Phylogenomic analysis independently supported this placement: FM48a grouped with *O. ostreae* CCUG 60525 with SH-aLRT/UFBoot support of 100/100, while *O. profunda* and other *Oceanisphaera* species formed separate lineages (Fig. 5A). GToTree recovered 72 of 74 bacterial marker genes from FM48a, corresponding to 97.3% marker recovery.

**FIG 5.**
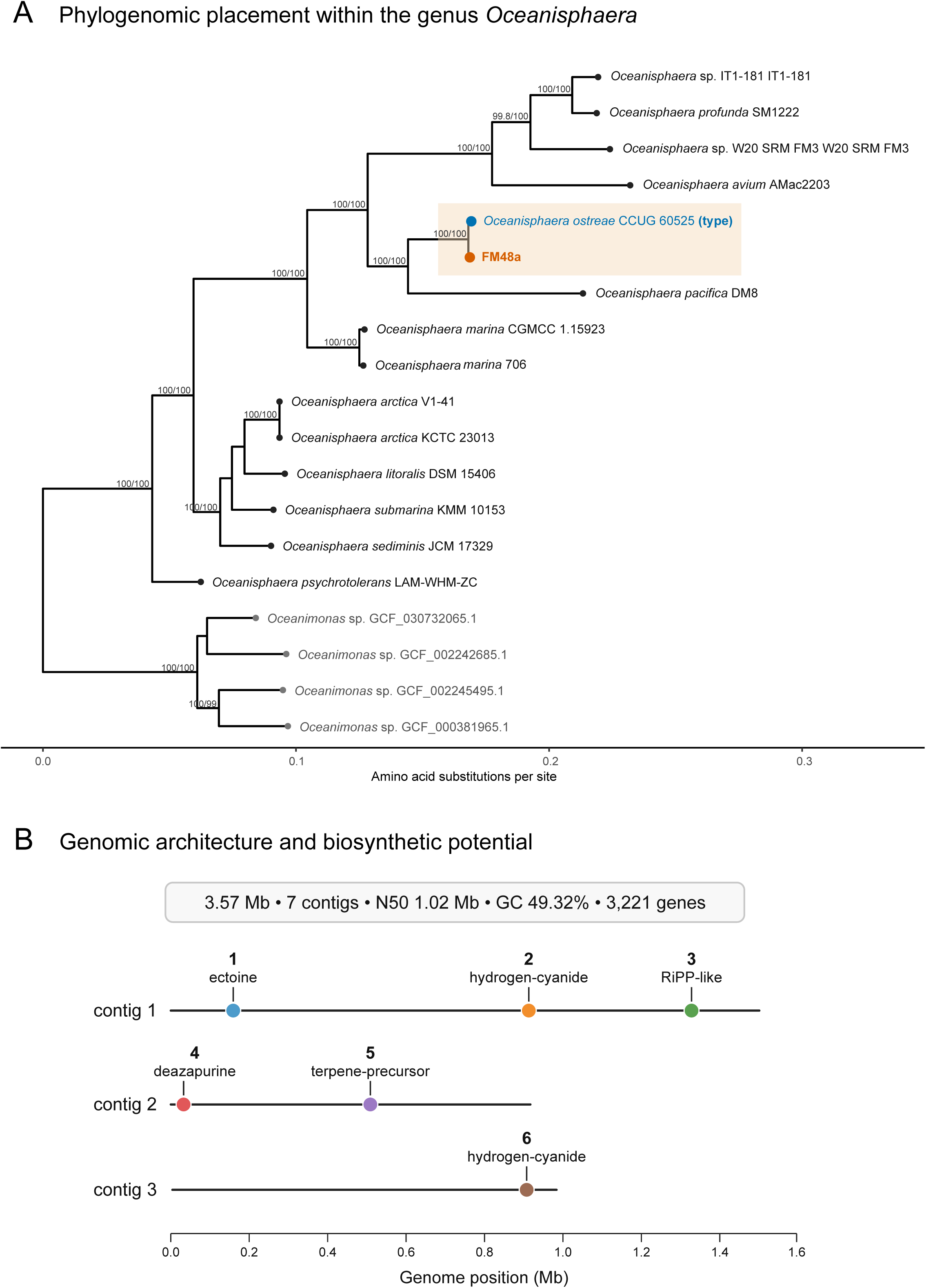
Phylogenomic placement, genomic biosynthetic potential and pilot bioactivity of *Oceanisphaera ostreae* FM48a. (A) Maximum-likelihood concatenated-marker phylogeny of *Oceanisphaera* and reference taxa; FM48a and the *O. ostreae* CCUG 60525 type-derived strain are highlighted. Node labels indicate SH-aLRT/ultrafast bootstrap support where shown. (B) FM48a assembly summary (3.57 Mb; seven contigs; N50 1.02 Mb; 49.32% GC; 3,221 genes) and approximate positions of six antiSMASH-predicted secondary-metabolite-related regions on contigs 1–3: ectoine, two hydrogen-cyanide-associated regions, RiPP-like, deazapurine and terpene precursor.

### Genome mining, metabolite profile, and preliminary bioactivity of FM48a

Genome mining with antiSMASH under relaxed strictness predicted six secondary-metabolite-related regions across the first three contigs (Fig. 5B). These comprised a predicted ectoine biosynthetic gene cluster (BGC) with high similarity to a known ectoine cluster, two hydrogen-cyanide-associated gene regions, one RiPP-like BGC, one deazapurine and one terpene-precursor related gene/cluster. No secondary-metabolite regions were reported on contigs 4-7.

LC–UV–MS analysis of the FM48a extract revealed a diverse metabolite profile, with multiple m/z features detected in both positive and negative ionization modes (Fig. 6A–D). Preliminary spectral comparisons suggested the presence of potentially bioactive compounds, including metabolites related to known microbial products such as lumichrome, a riboflavin-derived compound often found in marine bacteria that can be generated through microbial metabolism or photodegradation and has been associated with microbial signaling and interspecies interactions (18, 19).

**FIG 6.**
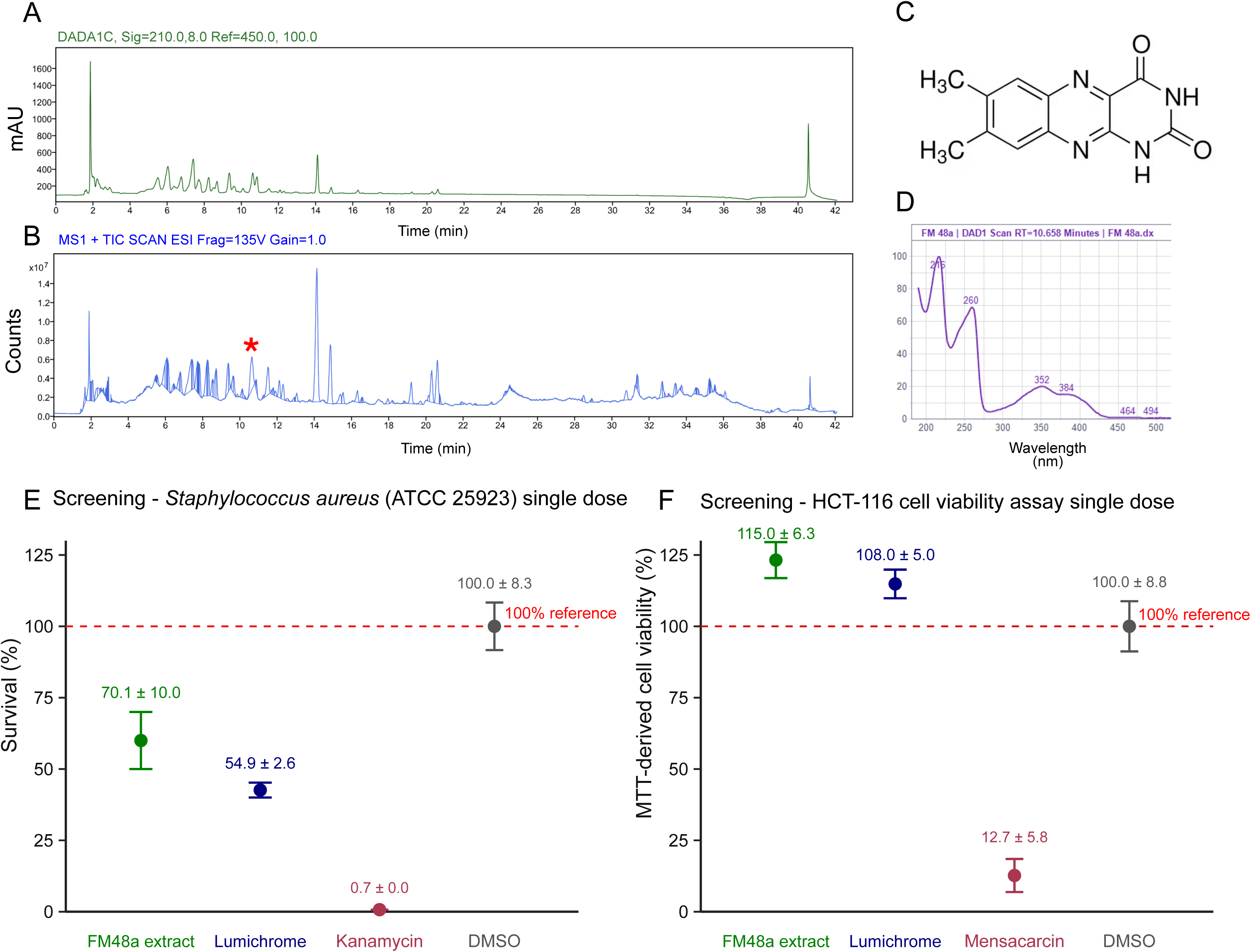
Chemical analysis and bioactivity. (A) 210-nm LCMS UV trace and (B) positive mode, electrospray (ESI) total ion chromatogram (TIC) of FM48a extract. Red asterisk indicates lumichrome, structure shown in (C) with a standard verified retention time of 10.7 minutes and characteristic UV absorption (D) and m/z value of 143 (M+H)^+^. (E) *Staphylococcus aureus* ATCC 25923 OD600-based relative growth/survival readouts after FM48a extract (125 μg/mL), lumichrome (10 μM), kanamycin (100 μg/mL, positive control), and DMSO (negative control) treatment (mean ± SEM). Dashed lines denote the non-active, 100% survival reference. (F) HCT-116, human colon carcinoma, MTT based cell viability readout after treatment with FM48a extract (10 μg/mL), lumichrome (10 μM), mensacarcin (10 μM, positive control), and DMSO (negative control) (mean ± SEM). Dashed lines denote the non-active, 100% survival reference.

In preliminary bioactivity screening, the FM48a extract caused 70.1 ± 10.0% survival of *S. aureus* ATCC 25923, while lumichrome reduced cell survival to 54.9 ± 2.6% under the tested assay conditions (Fig. 6E). In the HCT-116 MTT assay, mammalian cell survival was 115 ± 6.3% for the FM48a extract and 108 ± 5.0% for lumichrome, not affecting cell viability (Fig. 6F). These data indicate antibacterial activity in the screening assay but no evidence of HCT-116 cytotoxicity at the tested conditions.

## DISCUSSION

### Host-associated bacterial diversity varies among compartments but with high within-compartment variability

The bacterial microbiome of *H. antarcticus* is structured across anatomical compartments, while retaining considerable variability among individuals. The gut, peduncle and tentacles all harboured diverse bacterial assemblages, but alpha-diversity patterns were not uniform across the body. Gut samples showed the broadest spread in observed ASV richness and Fisher’s alpha, including several high-richness samples, whereas peduncle samples were generally less variable and tentacle samples showed intermediate to high diversity depending on the metric. This pattern suggests that the gut represents a particularly dynamic microbial compartment.

The high variability observed in gut samples is consistent with the feeding ecology of stauromedusae. *Haliclystus antarcticus* and related stalked jellyfishes are benthic predators that consume both planktonic and benthic prey, including small crustaceans and other invertebrates (16, 17). The gut microbiome may therefore integrate bacteria derived from prey, suspended particles, surrounding seawater and transient ingested material. This would explain why richness-based metrics showed stronger compartmental differences than Shannon or inverse Simpson diversity: gut samples may accumulate more rare or low-abundance taxa without necessarily changing the dominance structure of the community to the same extent.

At the same time, the UpSet analysis showed that the microbiome is not completely compartment-specific. The presence of 47 ASVs shared across gut, peduncle and tentacle samples indicates a recurrent host-associated bacterial pool distributed across the animal body. This shared component is important because it suggests that the *H. antarcticus* microbiome appears to include both broadly distributed host-associated taxa and compartment-enriched components. This organization is consistent with the unusual staurozoan body plan, in which a single attached adult combines polyp-like and medusa-like anatomical features (12).

### Taxonomic composition and bacterial community structure differ by site and compartment

The taxonomic composition of *H. antarcticus*-associated bacterial communities was dominated by bacterial groups commonly encountered in marine and cold-water systems, including *Paracoccaceae, Flavobacteriaceae, Methylophilaceae, Stappiaceae, Spongiibacteraceae* and *Moritellaceae.* Several abundant genera, such as *Polaribacter*, *Sulfitobacter*, *Methylotenera*, *Sinobacterium*, *Colwellia*, *Aliivibrio* and members of the Roseobacter clade, are consistent with bacterial groups frequently associated with seawater, particles, algal surfaces, organic matter degradation and animal-associated habitats. The presence of many unclassified genus-level assignments also highlights the limited representation of Antarctic host-associated bacteria in current reference databases.

The ordination analyses revealed differences in bacterial community composition associated with both host anatomical compartment and sampling site. Bray–Curtis dissimilarity showed strong effects of site and compartment, whereas weighted UniFrac indicated that compartment explained a substantial fraction of phylogenetic community variation. This suggests that compartmental structuring is not only a shift in the relative abundance of closely related taxa, but also involves phylogenetically distinct bacterial lineages. The separation of tentacle samples in ordination space is particularly notable and supports the idea that the tentacular region forms a distinct host–environment interface.

The significant effect of sampling site suggests that geographic context may contribute to variation in the *H. antarcticus*-associated bacterial communities. Although Byers Bay and Risopatron are geographically distinct, the environmental factors underlying the observed differences remain unclear. Local conditions, such as hydrodynamics, substrate-associated microbial communities, prey availability, and surrounding seawater bacterial assemblages, could potentially contribute to these patterns. This is especially relevant for a benthic, substrate-attached animal whose body regions experience different degrees of exposure to prey, seawater and substrate-associated bacteria.

The genus-level heatmap further supports this interpretation. Some taxa were widespread across samples, whereas others showed clear enrichment in subsets of compartments or sites. For example, genera such as *Methylotenera*, *Sinobacterium*, *Polaribacter* and *Sulfitobacter* contributed prominently across the dataset, while *Colwellia*, *Zobellia*, *Aurantivirga*, *Rubritalea*, *Maribacter* and *Aliivibrio* showed stronger enrichment in particular sample groups. These patterns suggest that microbiome differentiation is driven by consistent shifts in dominant bacterial taxa rather than by broad diversity changes alone.

### Compartment-specific bacterial associations

Differential-abundance and indicator value analyses provided additional evidence that bacterial communities associated with *H. antarcticus* vary among anatomical compartments. Gut and tentacle samples exhibited distinct taxonomic associations, suggesting that these anatomical regions represent different microbial microenvironments.

In the DESeq2 comparisons, gut samples showed enrichment of several bacterial ASVs relative to peduncle and tentacle samples. These included taxa assigned to genera commonly encountered in marine environments, such as *Colwellia*, *Maribacter*, *Aurantivirga*, and *Zobellia*. Indicator value analysis further identified gut-associated genera, including *Psychromonas*, *Granulosicoccus*, *Cocleimonas*, and *Marixanthomonas*. The enrichment of these taxa may reflect the distinctive ecological conditions of the gastrovascular cavity, where prey-derived material, ingested microorganisms, and organic substrates can accumulate. Such inputs could contribute to the greater variability in bacterial richness observed in gut samples. Given that stalked jellyfishes are active predators and consume both benthic and planktonic prey, the gut is likely a key compartment where environmental and trophic bacterial inputs are concentrated (16).

Tentacle-associated communities exhibited a different taxonomic pattern. DESeq2 identified several ASVs enriched in tentacles relative to the peduncle, while IndVal analysis identified *Polaribacter*, *Planktotalea*, *Perspicuibacter*, and *Peredibacter* as tentacle-associated indicator genera. The relatively high abundance of *Polaribacter* in tentacle samples is particularly notable. As tentacles represent a direct interface between the host and its surrounding environment, these associations may reflect repeated exposure to seawater, suspended particles, and prey-associated microorganisms.

In contrast, the peduncle did not exhibit significant single-compartment indicator genera under the applied criteria. This may indicate a less distinctive genus-level taxonomic signature compared with gut and tentacle samples. Given its role in substrate attachment, the peduncle may be exposed to a mixture of host-associated bacteria and microorganisms originating from surrounding benthic surfaces. In staurozoans, the peduncle and attachment-associated structures are central to the benthic lifestyle, while rhopalioids and adhesive organs have complex anatomical and neuromuscular organization (11).

The co-abundance analysis revealed groups of bacterial genera with positively and negatively correlated abundance profiles. These patterns may reflect shared responses to anatomical compartment, geographic location, or environmental conditions. For example, the positive correlations observed among *Polaribacter*, *Planktotalea*, *Sulfitobacter*, and related genera may be associated with similar distributions across host compartments. However, because these relationships were inferred from compositional 16S rRNA gene data, they should be interpreted as statistical associations rather than evidence of direct microbial interactions.

Overall, the combination of differential-abundance, indicator value, and co-abundance analyses supports the presence of anatomically structured bacterial communities in *H. antarcticus*. The distinct taxonomic associations observed in gut and tentacle samples suggest that different regions of the staurozoan body provide contrasting microbial habitats, potentially influenced by feeding activity, environmental exposure, and local ecological conditions.

### Broader implications for staurozoan microbiomes

Staurozoa remain one of the least explored cnidarian groups from a microbiome perspective. The first microbiome study of a stalked jellyfish, focused on *H. auricula* from the Barents Sea, showed that staurozoans can host diverse bacterial communities dominated by cold-adapted or psychrotolerant taxa, with *Colwellia* and *Marinomonas* among the dominant genera (7). Our study expands this perspective by showing that an Antarctic species, *H. antarcticus*, hosts a compartmentally structured bacterial microbiome.

This is important because staurozoans have a distinct body organization and ecology compared with more commonly studied cnidarians. They are benthic, attached, cryptic predators with limited dispersal and no free-swimming adult medusa stage. Reviews of staurozoan natural history emphasize their attachment to algae, rocks and other substrates, their shallow-water occurrence, and their feeding on small invertebrates such as amphipods and copepods (20). These features create multiple microbial interfaces: the peduncle with substrate biofilms, the tentacles with seawater and prey, and the gut with ingested material. The compartment-level microbiome structure observed here fits this ecological organization.

Overall, our results suggest that staurozoans are promising systems for studying how host anatomy, benthic lifestyle and polar coastal environments jointly shape animal-associated microbiomes. Future work integrating paired seawater, substrate and prey microbiomes would help clarify the relative contribution of environmental source pools and host filtering, while metagenomic or microscopy-based approaches could identify whether dominant bacterial taxa are transient associates or functionally active members of the host microbiome.

### Cnidarian microbiome context

The compartment-level structure observed in *H. antarcticus* fits a broader pattern emerging from cnidarian microbiome studies: bacterial communities are often shaped by host life stage, anatomy, symbiotic state and environment rather than simply mirroring surrounding seawater. In scyphozoan jellyfish, microbiomes can differ among life stages and include recurrent bacterial cores (4, 21). Recent work on jellyfish development further suggests that cnidarian-associated bacteria can influence host phenotype and life-history transitions (6, 22). In Aurelia coerulea von Lendenfeld, 1884, microbiota manipulation affected strobilation, nematocyte development and feeding-related traits, indicating that bacterial communities can contribute to the regulation of jellyfish development rather than acting only as passive colonizers (23, 24). In *Cassiopea xamachana* Bigelow, 1892, bacterial communities vary across anatomy, development and geography, with internal communities reported as distinct from external surfaces and the surrounding environment (6, 25). These findings are consistent with our observation that *H. antarcticus* compartments harbour related but distinct bacterial assemblages.

Across cnidarians, microbiomes appear to be shaped by host biological state, local microhabitat and environmental exposure. Our results therefore extend cnidarian microbiome research into Staurozoa and suggest that fine-scale anatomical partitioning may be a general but underexplored feature of cnidarian-associated bacterial communities.

### Characterization of *Oceanisphaera ostreae*: biosynthetic potential, metabolite profiling, and biological activity

The isolation and characterization of *O. ostreae* FM48a from the gut of *H. antarcticus* provide a cultivation-based perspective that complements the culture-independent characterization of the host-associated bacterial community. Genome mining of FM48a identified six putative biosynthetic gene clusters, suggesting the potential to produce metabolites involved in physiological adaptation and ecological interactions. Among these, the predicted ectoine biosynthetic region is particularly relevant in the context of Antarctic marine environments, as ectoine is a compatible solute associated with bacterial responses to osmotic and other environmental stresses. Its predicted biosynthetic capacity may therefore reflect physiological adaptations that facilitate survival under variable environmental conditions. However, ectoine production and its contribution to stress tolerance remain to be experimentally confirmed. Additional regions annotated as RiPP-like, deazapurine, and terpene-precursor pathways suggest further metabolic potential, and might encode for compounds involved in microbial interactions or environmental responses. Two regions associated with hydrogen cyanide biosynthesis were also predicted, although their presence alone does not establish hydrogen cyanide production or any corresponding biological activity. Collectively, these genomic findings highlight the biosynthetic potential of FM48a, while emphasizing that computational predictions require complementary chemical and functional validation. It is also noteworthy that genome mining techniques are still limited among marine Gram-negative bacteria. We recently published novel benzothiazine and benzothiazole from a marine *Croceibacter atlanticus* strain with analgesics activity, which could not be predicted from its genome (26). LC–UV–MS analysis of the FM48a extract revealed a complex chemical profile containing multiple chromatographic features detected in positive and negative ionization modes (Fig. 6). Lumichrome, a photodegradation product of riboflavin, is of potential biological interest because flavin-related compounds can participate in microbial redox processes and ecological interactions.

Preliminary biological screening provided additional evidence that FM48a produces extractable material capable of influencing bacterial growth-related measurements. When tested against *Staphylococcus aureus* ATCC 25923, the FM48a extract resulted in a mean relative OD600-based survival readout of 70.1 ± 10.0%, compared with 54.9 ± 2.6% for the separately tested lumichrome standard. These observations suggest a potential inhibitory effect under the experimental conditions, although they do not establish bactericidal activity or identify the compound responsible. In contrast, screening against HCT-116 human colorectal carcinoma cells did not affect mammalian cell viability under the conditions examined. MTT-based viability measurements were 115 ± 6.3% for the FM48a crude extract and 108 ± 5.0% for lumichrome relative to the assay control, suggesting that neither treatment substantially reduced the metabolic viability signal at the concentrations tested.

Taken together, the cultivation, genomic and chemical characterization of *O. ostreae* FM48a expands the functional perspective of the *H. antarcticus*-associated microbiome beyond taxonomic community profiling. The identification of predicted biosynthetic pathways and the preliminary antibacterial screening results suggest that cultivable gut-associated bacteria may represent a source of metabolically diverse compounds with potential ecological and biotechnological relevance.

## CONCLUSIONS

This study provides the first compartment-resolved description of the bacterial microbiome associated with the Antarctic stalked jellyfish *H. antarcticus*. Across host-associated samples from Byers Bay and Risopatron, we identified a diverse bacterial assemblage composed of both shared and compartment-enriched ASVs.

The microbiome was structured by host anatomy: gut, peduncle and tentacle samples differed in alpha diversity, taxonomic composition, ordination space and marker taxa. Gut samples showed the highest variability, consistent with exposure to prey-associated and ingested microbial inputs, while tentacles harboured distinct indicator taxa such as *Polaribacter* and *Planktotalea*.

The isolation of *O. ostreae* from the gut further highlighted the metabolic and biosynthetic potential of host-associated bacteria. Genomic analyses revealed several putative biosynthetic pathways, including those associated with ectoine production and secondary metabolism, while metabolite profiling identified chemically diverse features, including lumichrome. Preliminary antibacterial activity against *S. aureus* further suggests that gut-associated bacteria may produce bioactive metabolites with potential ecological and biotechnological relevance.

Together, these findings show that *H. antarcticus* hosts a compartmentally structured bacterial community shaped by both host anatomy and geographic context. More broadly, the study expands cnidarian microbiome research to an understudied Antarctic staurozoan and highlights the biosynthetic potential and possible ecological relevance of their associated bacteria.

## MATERIAL AND METHODS

### Sample collection and compartment dissection

A total of 65 individuals of *H. antarcticus* were collected in January 2024 from two sites along the coasts of the South Shetland Islands: Byers Bay (62°38.34′S, 60°36.9′W), and Risopatron (62°22.65′S, 59°42.08′W), at a depth of about 1 m (Fig. 1A-D).

Specimens were found on coarse gravel substrata and attached to macroalgae, including kelp, and were carefully detached to preserve the peduncle and associated tissues. After collection, individuals were rinsed with sterile 0.2 µm-filtered artificial seawater to reduce loosely associated microorganisms. Each individual was dissected under sterile conditions into three anatomical compartments: gut, peduncle, and tentacles (Fig. 1E). Dissection tools were sterilized between samples to minimize cross-contamination among individuals and compartments. Samples were stored at −20°C in RNA later until DNA extraction.

### Amplicon gene region sequencing

DNA was extracted from gut, tentacle and peduncle tissues of 45 individuals from Byers Bay and 20 individuals from Risopatron using DNeasy PowerSoil kit (QIAGEN, CA, USA) following manufacturer’s instructions with few modifications (vortexing extended to 20 min, and final elution volume reduced to 25 µL). Three control samples were processed in parallel with all samples to identify and account for any contaminant ASVs introduced during extraction, amplification, or sequencing. The 16S rRNA gene V4 hypervariable region was amplified using primer set 515F (Parada) (5′-GTGYCAGCMGCCGCGGTAA-3′) and 806R (Apprill) (5′-GGACTACNVGGGTWTCTAAT-3′) (27, 28). This primer set was selected because it is widely used for marine microbial community profiling and cnidarian-associated microbiome studies. (4, 5). PCR amplification was performed using the following conditions: initial denaturation at 95°C for 3 min, followed by 30 cycles of denaturation at 95°C for 30 s, annealing at 55°C for 30 s, and extension at 72°C for 30 s, with a final extension at 72°C for 5 min. Amplicon concentrations were confirmed with a Qubit 3 fluorometer (FisherSci) and quality was checked on a spectrophotometer (BioTek Take 3 Plate, Agilent) using the 260/280 nm ratio. DNA concentrations were normalized prior to sequencing. Libraries were sequenced at Genewiz (Azenta) on an Illumina MiSeq platform (2 × 250 bp).

### Microbiome analysis

Sequences were filtered and trimmed using the DADA2 R-package (v1.22 (29)) in RStudio (2024.04.2 (30)), including quality filtering, error rate learning, denoising, merging and chimera removal. Sequences were processed according to the DADA2 SOP (https://benjjneb.github.io/dada2/tutorial.html) using standard filtering (maxN = 0, maxEE = 1,1), denoising, merging, chimera and singletons removal, and naïve Bayesian taxonomic classification. To assess and minimize potential contamination at different stages of the workflow, negative controls were included, comprising water blanks (autoclaved artificial seawater filtered through a 0.2-µm membrane), extraction blanks, primer blanks, and PCR blanks. Potential contaminant amplicon sequence variants (ASVs) were identified using the frequency-based method implemented in the decontam R package (v1.24.0 (31)), with a classification threshold of 0.1. A total of 33 ASVs were identified as potential contaminants and removed before downstream analyses.

Taxonomic annotation of sequences was performed with the Silva database (v. 138.2 SSU (32)). ASVs not classified to the Kingdom ‘Bacteria’, or assigned as ‘Chloroplast’ were removed. Sequence data and associated metadata were imported into phyloseq R-package (v1.50.0 (33)) for downstream analysis.

### Statistical analysis

All statistical analyses were conducted in RStudio (2024.04.2 (30)). For microbiome analyses, amplicon sequence variant (ASV) tables and metadata were imported into phyloseq (v1.50.0 (33)) for data handling and visualization, using rstatix (v0.7.2 (34)) and ggplot2 (v4.0.0 (35)).

Alpha diversity was calculated at the ASV level using four metrics: observed ASV richness, Fisher’s alpha, Shannon diversity and inverse Simpson diversity (34, 36–38). These metrics were selected to capture complementary aspects of within-sample diversity, including richness, richness-adjusted diversity and evenness-sensitive diversity. Alpha diversity was analysed using linear mixed-effects models with compartment and site as fixed effects and individual identity as a random intercept, accounting for the repeated sampling of multiple compartments from the same host individual. Models were fitted using lme4 and lmerTest (v1.1.37 (39)). Pairwise comparisons among compartments were performed using estimated marginal means with Benjamini–Hochberg adjustment for multiple testing.

Shared ASVs among host compartments were examined using an UpSet-style intersection analysis using ComplexUpset (v1.3.3 (40)). ASVs were considered present in a compartment if detected in at least one-third of samples from that compartment. Compartment-level ASV sets were then compared across gut, peduncle and tentacle samples to identify ASVs shared across all compartments and ASVs preferentially detected in one or two compartments. For taxonomic composition analyses, ASVs were agglomerated independently at family and genus levels using phyloseq. Counts were transformed to relative abundance within each sample. For visualization, the most abundant families and genera were retained, and lower-abundance taxa were grouped as “Other”.

Community dissimilarity among samples was assessed using Bray–Curtis dissimilarity and weighted UniFrac distance. Bray–Curtis dissimilarity was calculated from ASV count tables. Weighted UniFrac distances were calculated using a phylogenetic tree associated with the ASV table and incorporated relative abundance and phylogenetic relatedness among ASVs. Differences in bacterial community composition were tested using permutational multivariate analysis of variance (PERMANOVA) with 9,999 permutations using vegan::adonis2 (v2.6.10 (41)). Models included site and compartment as explanatory variables. Because multiple compartments were sampled from the same individual, compartment effects were also evaluated using permutation schemes constrained by individual identity. Homogeneity of multivariate dispersion was assessed using betadisper and permutation tests.

To visualize dominant genus-level patterns across samples, ASVs were agglomerated at the genus level and transformed to relative abundances. The most abundant genera were retained for visualization, and their relative abundances were log10-transformed.

Differential abundance among host anatomical compartments was assessed at the ASV level using DESeq2 (v1.46.0 (42)) on unrarefied count data. The statistical model included sampling site and anatomical compartment as explanatory variables to account for geographic variation when testing for compartment-associated differences. Pairwise comparisons were performed between gut and peduncle, gut and tentacle, and peduncle and tentacle samples. ASVs were considered significantly differentially abundant when the Benjamini–Hochberg-adjusted *P* value was ≤ 0.05. Log2 fold changes were used to describe the direction and magnitude of differences between compartments.

Indicator taxa associated with host anatomical compartments were identified at the genus level using the indicspecies R package. Indicator value analysis was performed using the multipatt function with the IndVal.g statistic and 999 permutations. The analysis was restricted to single-compartment associations (duleg = TRUE), and genera with *P* ≤ 0.05 were considered significant compartment indicators.

Exploratory genus-level co-abundance patterns were assessed using Spearman rank correlations calculated from centred log-ratio (CLR)-transformed relative abundance profiles. Genera were retained if they occurred in at least 20% of host-associated samples and had a mean relative abundance ≥ 0.1%. Among genera meeting these criteria, the 50 most abundant were selected for analysis. A small pseudocount was added before CLR transformation to accommodate zero values. Pairwise Spearman correlation coefficients were calculated among retained genera, and *P* values were adjusted for multiple comparisons using the Benjamini–Hochberg procedure. The resulting correlation matrix was visualized as a hierarchically clustered heatmap to explore patterns of positive and negative co-abundance among bacterial genera.

### Isolation, genome characterization of *Oceanisphaera ostreae*

A bacterial strain (FM48a) was isolated from intestinal material of an *H. antarcticus* specimen collected at Risopatron. A 200-µL aliquot was plated directly on SYP-SW (Starch Yeast Peptone SeaWater) media. FM 48a was grown by using a sterile loop to inoculate a single colony into 50 mL of autoclave-sterilized SYP-SW medium (10 g/L starch, 4 g/L yeast, 2 g/L peptone, 33.3 g/L instant ocean, MQ water). Three starter cultures were incubated for two weeks at room temperature (22-23°C), orbitally shaking at 150 rotations per minute. The cultures were clean streaked onto SYP-SW plates (same media plus 15 g/L agar) to confirm there were no signs of contamination. Using a sterile serological pipette, 25 mL of each culture was then used to inoculate three 1L cultures of SYP-SW media. These cultures were grown for six weeks under the same temperature and shaking conditions as the 50 mL cultures. The cultures were combined and extracted by liquid-liquid partition using equal volumes of ethyl acetate and then concentrated into an extract using rotary evaporation.

For DNA extraction, 10 mL of turbid SYP-SW culture of FM 48a was transferred to a 15 mL falcon tube and centrifuged at 3000 RPM for 10 minutes. Most of the liquid media was decanted from the cell pellet and roughly 3.0 mL of remaining media was used to resuspend the pellet and divide the contents into two 2.0 mL Eppendorf tubes. These tubes were spun down at 3000 rpm for 10 minutes and the remaining media decanted. The pellet was then frozen at -80°C overnight and then the cells lysed using a mortar and pestle chilled to -20°C in combination with periodic sonication. The crushed cells were then suspended in TE buffer supplemented with lysozyme (20 mg/mL) and incubated for 30 minutes at 37°C. To this mixture, 3 µL of proteinase K, and 55 µL of SDS (10% w/v) were added and the new mixture was incubated for 120 minutes at 55°C. After cooling to room temperature, the mixture was extracted using phenol:chloroform:isoamyl alcohol (25:24:1) and then centrifuged at 14,000 rpm for 10 minutes. The aqueous layer was transferred into a sterile Eppendorf tube and the DNA was precipitated by adding 3M sodium acetate (10% v/v) and then 400 µL of cold isopropanol. The DNA was then extracted using centrifugation on a Qiagen spin column and C6 elution buffer. DNA purity was assessed using 260/280 and 260/230 ratios on a NanoDrop and the final DNA concentration was measured by Qubit to be 59 ng/µL. The sample was then submitted to commercial sequencing (Plasmidsaurus) for long-read, nanopore sequencing. The *O. ostreae* FM48a isolate was registered in the NCBI BioSample database under accession number SAMN64870342.

Whole-genome sequencing, assembly and annotation were performed for this strain. Assembly statistics were calculated from the final genome FASTA. Genome-based taxonomic placement was assessed with FastANI v1.33 (43, 44) against 30 publicly available *Oceanisphaera* assemblies from NCBI, including genomes derived from type material.

Reciprocal ANI comparisons were performed, and the mean reciprocal ANI and minimum reciprocal alignment fraction were recorded. ANI ≥95% together with alignment fraction ≥0.50 was used as a species-level screening criterion rather than as an automatic nomenclatural assignment.

Phylogenomic placement was evaluated with GToTree v1.8.19 (45) using the bacterial single-copy marker HMM set (74 targets). The *Oceanisphaera* reference set was deduplicated to one representative per biological strain for interpretation. Marker proteins were aligned, trimmed and concatenated, and a maximum-likelihood tree was inferred with IQ-TREE 3.1.3 (46). ModelFinder (47) selected Q.INSECT+F+R4, and branch support was evaluated with 1,000 ultrafast-bootstrap replicates (48) and 1,000 SH-aLRT replicates (49). *Oceanimonas* and *Dongshaea* genomes were included as outgroups in the initial inference; redundant GenBank/RefSeq representations were pruned for the final visualization.

The genome annotation contained seven 16S rRNA gene copies. The 515F-806R V4 region was extracted from each copy and deduplicated. All seven copies carried the same 253-bp V4 sequence, which was compared directly with DADA2 ASV sequences from the raw sequence table before prevalence filtering and with the raw *H. antarcticus* host-only table. An isolate-linked amplicon match was defined as 100% identity across the complete 253-bp FM48a V4 sequence, allowing only short terminal overhangs in the merged amplicon.

### Genome mining, LC-MS profiling and preliminary bioactivity screening

Biosynthetic and secondary-metabolite-related genomic regions were screened with antiSMASH 8.0 with the detection strictness set to relaxed (50). FM48a organic extracts were analyzed by low-resolution LC-MS in positive and negative ionization modes with diode-array UV detection at 210, 254, 280 and 360 nm. Features near retention times 10.5 and 10.8 min were examined in detail, and the UV spectrum of the ∼10.7-min feature was compared with an internal UV spectral library and identified by UV and mass to be lumichrome, often found in marine *Gammaproteobacteria* (51). Low-resolution ESI-MS analysis was performed with an Agilent 1200 series LC system and an Agilent Pro iQ Plus MSD, using a Phenomenex Kinetex C18 5 µm, 150 x 4.6 mm column. Each injection contained 10 µL of sample over a 33 minute gradient from 10% to 100% ACN, 90%-0% MilliQ water, both supplemented with 0.05% (v/v) LCMS grade formic acid, with a flow rate of 0.8 mL/min. MS conditions for both positive and negative mode acquisition were as follows: 350°C drying gas temperature, 13.0 L/min drying gas flow rate, 35 nebulizer psi, 3000 V capillary voltage, 250°C sheath gas temperature, 11.0 L/min sheath gas flow rate, 1500 V nozzle voltage.

Preliminary bioactivity of the FM48a organic extract and the identified bacterial metabolite lumichrome was evaluated against *Staphylococcus aureus* ATCC 25923 using OD600 as a growth/survival readout and against HCT-116 colon cancer cells using an MTT colorimetric viability assay. For the *S. aureus* assay, the final concentration of samples was 125 µg/mL in MHII broth, with a kanamycin positive control, DMSO negative control, and run in technical replicates incubated for 18 hours at 37°C without shaking. Changes to optical density at 600 nm were measured relative to uninoculated media. For the colorimetric MTT assay using HCT-116 colon cancer cells, the final concentration of samples was 10 µM, mensacarcin was used as the positive control, DMSO was used as the negative control, and samples were assayed in technical duplicate. HCT-116 cells were seeded into a 96-well plate at a density of 7000 cells per well and allowed to adhere for 24 hours in a humidified incubator with 5% CO2 at 37°C. Test samples solubilized in DMSO were added to the seeded wells and incubated for a period of 46 hours. Then, MTT solubilized in PBS was added to each well at a concentration of 0.5 mg/mL and after two hours the DMEM was removed from the wells and the remaining cells and formazan crystals resuspended in 50µL of DMSO and absorbance measured at 550 nm. Metabolic activity of DMSO-treated cells was considered 100% viability and samples that reduced metabolic activity to 60% or less were considered anti-proliferative.

## ACKNOWLEDGMENTS

The authors gratefully acknowledge the Instituto Antártico Chileno (INACH) for the opportunity to participate in the 60th Antarctic Scientific Expedition (ECA 60) and for the logistical support provided during fieldwork in Antarctica. We thank Julia Sutton for her assistance with DNA isolation, Julia was supported by an American Society of Pharmacognosy (ASP) summer fellowship.

## DATA AVAILABILITY STATEMENT

Sequence data generated in this study are available through the NCBI BioProject under accession number PRJNA1558909. The *Oceanisphaera ostreae* FM48a isolate is registered under NCBI BioSample accession number SAMN64870342. The FM48a draft genome has been submitted to NCBI, and its genome accession number will be included following assignment. The complete analysis code is available on GitHub at https://github.com/federicamontesanto/Haliclystus_microbiome.

## REFERENCES

1. Bourne DG, Morrow KM, Webster NS. 2016. Insights into the coral microbiome: underpinning the health and resilience of reef ecosystems. Annu Rev Microbiol 70:317–340.

2. Rädecker N, Pogoreutz C, Voolstra CR, Wiedenmann J, Wild C. 2015. Nitrogen cycling in corals: the key to understanding holobiont functioning? Trends Microbiol 23:490–497.

3. Voolstra CR, Raina J-B, Dörr M, Cárdenas A, Pogoreutz C, Silveira CB, Mohamed AR, Bourne DG, Luo H, Amin SA, Peixoto RS. 2024. The coral microbiome in sickness, in health and in a changing world. Nat Rev Microbiol 22:460–475.

4. McCauley M, Goulet TL, Jackson CR, Loesgen S. 2023. Systematic review of cnidarian microbiomes reveals insights into the structure, specificity, and fidelity of marine associations. Nat Commun 14:4899.

5. McCauley M, Montesanto F, Bedgood SA, Miner C, Plichon K, Weis VM, Loesgen S. 2025. Manipulation of the Symbiodiniaceae microbiome confers multigenerational impacts on symbioses and reproductive ecology of its *Exaiptasia diaphana* host. ISME J 19:wraf189.

6. Montesanto F, McCauley M, Bedgood SA, Miner C, Steinworth B, Sharp V, Ohdera AH, Oluokun A, Fowowe M, Oluokun O, Mechref Y, Xiang T, Medina M, Weis VM, Martindale MQ, Loesgen S. 2026. Cnidarian–algal partnerships structure bacterial communities during strobilation in *Cassiopea xamachana*. ISME Commun 6:ycag147.

7. Khabibulina V, Domracheva M, Sogomonyan K. 2025. First description of the Haliclystus auricula (Cnidaria: Staurozoa) jellyfish bacterial microbiome. Polar Biol 48:73.

8. Weiland-Bräuer N, Neulinger SC, Pinnow N, Künzel S, Baines JF, Schmitz RA. 2015. Composition of bacterial communities associated with *Aurelia aurita* changes with compartment, life stage, and population. Appl Environ Microbiol 81:6038–6052.

9. Kerwin AH, Ohdera A, Bier J, Goodman D, Mammone M, Sharp V, Echeandía A, Medina M. 2025. *Cassiopea xamachana* microbiome across anatomy, development, and geography. PLOS ONE 20:e0319944.

10. Murray AE, Rack FR, Zook R, Williams MJM, Higham ML, Broe M, Kaufmann RS, Daly M. 2016. Microbiome Composition and Diversity of the Ice-Dwelling Sea Anemone, *Edwardsiella andrillae*. Integr Comp Biol 56:542–555.

11. Domracheva M, Khabibulina V. 2024. Adhesive organs with complex neuromuscular system: Organization of rhopalioids in *Haliclystus auricula* James-Clark, 1863 (Cnidaria: Staurozoa). Acta Zool 105:403–418.

12. Miranda LS, Collins AG, Marques AC. 2013. Internal anatomy of *Haliclystus antarcticus* (Cnidaria, Staurozoa) with a discussion on histological features used in staurozoan taxonomy: Internal Anatomy Of Haliclystus Antarcticus. J Morphol 274:1365–1383.

13. Kos Kramar M, Tinta T, Lučić D, Malej A, Turk V. 2019. Bacteria associated with moon jellyfish during bloom and post-bloom periods in the Gulf of Trieste (northern Adriatic). PLOS ONE 14:e0198056.

14. Barak N, Brekhman V, Aharonovich D, Lotan T, Sher D. 2025. Jellyfish blooms through the microbial lens: temporal changes, cross-species and Jellyfish-water comparisons. Environ Microbiome 20:49.

15. Miranda LS, Morandini AC, Marques AC. 2009. Taxonomic review of *Haliclystus antarcticus* Pfeffer, 1889 (Stauromedusae, Staurozoa, Cnidaria), with remarks on the genus Haliclystus Clark, 1863. Polar Biol 32:1507–1519.

16. Davenport J. 1998. Note on the Trophic Relationships of the Stauromedusa *Haliclystus antarcticus* from Subantarctic South Georgia. J Mar Biol Assoc U K 78:663–664.

17. Zagal CJ. 2004. Diet of the stauromedusa *Haliclystus auricula* from southern Chile. J Mar Biol Assoc U K 84:337–340.

18. Yamamoto K, Asano Y. 2015. Efficient Production of Lumichrome by Microbacterium sp. Strain TPU 3598. Appl Environ Microbiol 81:7360–7367.

19. Rajamani S, Bauer WD, Robinson JB, Farrow JM, Pesci EC, Teplitski M, Gao M, Sayre RT, Phillips DA. 2008. The Vitamin Riboflavin and Its Derivative Lumichrome Activate the LasR Bacterial Quorum-Sensing Receptor. Mol Plant-Microbe Interactions® 21:1184–1192.

20. Miranda LS, Mills CE, Hirano YM, Collins AG, Marques AC. 2018. A review of the global diversity and natural history of stalked jellyfishes (Cnidaria, Staurozoa). Mar Biodivers 48:1695–1714.

21. Röthig T, Puntin G, Wong JCY, Burian A, McLeod W, Baker DM. 2021. Holobiont nitrogen control and its potential for eutrophication resistance in an obligate photosymbiotic jellyfish. Microbiome 9:127.

22. Carabantes N, Cerqueda-García D, García-Maldonado JQ, Thomé PE. 2022. Changes in the bacterial community associated with experimental symbiont loss in the mucus layer of *Cassiopea xamachana* jellyfish. Front Mar Sci 9:879184.

23. Peng S, Ye L, Li Y, Wang F, Sun T, Wang L, Hao W, Zhao J, Dong Z. 2023. Microbiota regulates life-cycle transition and nematocyte dynamics in jellyfish. iScience 26:108444.

24. Peng S, Ye L, Li Y, Wang F, Sun T, Wang L, Zhao J, Dong Z. 2024. Metagenomic insights into jellyfish-associated microbiome dynamics during strobilation. ISME Commun 4:ycae036.

25. Kerwin AH, Ohdera A, Bier J, Goodman D, Mammone M, Sharp V, Echeandía A, Medina M. 2025. *Cassiopea xamachana* microbiome across anatomy, development, and geography. PLoS ONE 20:e0319944.

26. Marshall EM, Zhu C, Mayhood L, Khalid Z, Yao H, Panieri G, Cediel-Becerra JDD, Chevrette MG, Strother JA, Loesgen S. 2026. Antinociceptive Benzothiazine and Benzothiazole Natural Products from Marine *Croceibacter atlanticus*. J Nat Prod 89:1265–1275.

27. Parada AE, Needham DM, Fuhrman JA. 2016. Every base matters: assessing small subunit rRNA primers for marine microbiomes with mock communities, time series and global field samples. Environ Microbiol 18:1403–1414.

28. Apprill A, McNally S, Parsons R, Weber L. 2015. Minor revision to V4 region SSU rRNA 806R gene primer greatly increases detection of SAR11 bacterioplankton. Aquat Microb Ecol 75:129–137.

29. Callahan BJ, McMurdie PJ, Rosen MJ, Han AW, Johnson AJA, Holmes SP. 2016. DADA2: High-resolution sample inference from Illumina amplicon data. Nat Methods 13:581–583.

30. Posit Team. 2025. *RStudio: Integrated Development Environment for R.* Boston, MA: Posit Software, PBC.

31. Davis NM, Proctor DM, Holmes SP, Relman DA, Callahan BJ. 2018. Simple statistical identification and removal of contaminant sequences in marker-gene and metagenomics data. Microbiome 6:226.

32. Quast C, Pruesse E, Yilmaz P, Gerken J, Schweer T, Yarza P, Peplies J, Glöckner FO. 2012. The SILVA ribosomal RNA gene database project: improved data processing and web-based tools. Nucleic Acids Res 41:D590–D596.

33. McMurdie PJ, Holmes S. 2013. phyloseq: An R-package for reproducible interactive analysis and graphics of microbiome census data. PLoS ONE 8:e61217.

34. Kassambara A. rstatix: pipe-friendly Framework for basic statistical tests. 2023. 0.7.2.

35. Wickham H. 2016. Getting Started with ggplot2, p. 11–31. *In* . Springer International Publishing, Cham.

36. Fisher RA, Corbet AS, Williams CB. 1943. The relation between the number of species and the number of individuals in a random sample of an animal population. J Anim Ecol 12:42.

37. Shannon CE. 1948. A mathematical theory of communication. Bell Syst Tech J 27:379–423.

38. Simpson EH. 1949. Measurement of diversity. Nature 163:688–688.

39. Bates D, Maechler M, Bolker B, Walker S. 2003. lme4: linear mixed-effects models using “Eigen” and S4.

40. Michał Krassowski, Arts M, Lagger C, Max. 2022. ComplexUpset: create Complex UpSet plots using “ggplot2” components. 1.3.3. 10.5281/ZENODO.3700590.

41. Dixon P. 2003. VEGAN, a package of R functions for community ecology. J Veg Sci 14:927–930.

42. Love MI, Huber W, Anders S. 2014. Moderated estimation of fold change and dispersion for RNA-seq data with DESeq2. Genome Biol 15:550.

43. Jain C, Rodriguez-R LM, Phillippy AM, Konstantinidis KT, Aluru S. 2018. High throughput ANI analysis of 90K prokaryotic genomes reveals clear species boundaries. Nat Commun 9:5114.

44. Richter M, Rosselló-Móra R. 2009. Shifting the genomic gold standard for the prokaryotic species definition. Proc Natl Acad Sci 106:19126–19131.

45. Lee MD. 2019. GToTree: a user-friendly workflow for phylogenomics. Bioinformatics 35:4162–4164.

46. Wong TKF, Ly-Trong N, Ren H, Demotte P, Baños H, Roger AJ, Susko E, Bielow C, De Maio N, Goldman N, Hahn MW, Dos Reis M, Vinh LS, Huttley G, Lanfear R, Minh BQ. 2026. IQ-TREE 3: phylogenomic inference software using complex evolutionary models. Mol Biol Evol 43:msag117.

47. Kalyaanamoorthy S, Minh BQ, Wong TKF, Von Haeseler A, Jermiin LS. 2017. ModelFinder: fast model selection for accurate phylogenetic estimates. Nat Methods 14:587–589.

48. Hoang DT, Chernomor O, Von Haeseler A, Minh BQ, Vinh LS. 2018. UFBoot2: Improving the Ultrafast Bootstrap Approximation. Mol Biol Evol 35:518–522.

49. Guindon S, Dufayard J-F, Lefort V, Anisimova M, Hordijk W, Gascuel O. 2010. New Algorithms and Methods to Estimate Maximum-Likelihood Phylogenies: Assessing the Performance of PhyML 3.0. Syst Biol 59:307–321.

50. Blin K, Shaw S, Vader L, Szenei J, Reitz ZL, Augustijn HE, Cediel-Becerra JDD, de Crécy-Lagard V, Koetsier RA, Williams SE, Cruz-Morales P, Wongwas S, Segurado Luchsinger AE, Biermann F, Korenskaia A, Zdouc MM, Meijer D, Terlouw BR, van der Hooft JJJ, Ziemert N, Helfrich EJN, Masschelein J, Corre C, Chevrette MG, van Wezel GP, Medema MH, Weber T. 2025. antiSMASH 8.0: extended gene cluster detection capabilities and analyses of chemistry, enzymology, and regulation. Nucleic Acids Res 53:W32–W38.

51. Rajamani S, Bauer WD, Robinson JB, Farrow JM, Pesci EC, Teplitski M, Gao M, Sayre RT, Phillips DA. 2008. The Vitamin Riboflavin and Its Derivative Lumichrome Activate the LasR Bacterial Quorum-Sensing Receptor. Mol Plant-Microbe Interactions® 21:1184–1192.

